# Wiskott-Aldrich Syndrome Protein Regulates Nucleolar Organization and Function in Innate Immune Response

**DOI:** 10.1101/2023.08.15.553387

**Authors:** Xuan Zhou, Baolei Yuan, Yeteng Tian, Juexiao Zhou, Mengge Wang, Ismail Shakir, Yingzi Zhang, Chongwei Bi, Bayan Mohammed Aljamal, Mais O. Hashem, Omar Imad Abuyousef, Firdous Mohammed Abdulwahab, Afshan Ali, Sarah Dunn, James J. Moresco, John R. Yates, Francesco Frassoni, Xin Gao, Fowzan S. Alkuraya, Juan Carlos Izpisua Belmonte, Mo Li

## Abstract

Wiskott-Aldrich syndrome (WAS) is a primary immunodeficiency disorder caused by the dysfunction of the WAS protein (WASP). Using an isogenic macrophage model derived from genome edited induced pluripotent stem cells we demonstrated that WASP functions in the nucleolus, which plays important roles in immune regulation. The absence of WASP resulted in smaller and misshapen nucleoli, decreased fibrillar center territory, and impaired ribosomal RNA (rRNA) transcription. The nucleolar and rRNA phenotypes were confirmed in WAS patient samples. Furthermore, WASP interacts with nucleolar proteins, including nucleophosmin 1 (NPM1) and fibrillarin (FBL). NPM1 deficiency is known to cause elevated cytokine expression following lipopolysaccharide (LPS) stimulation. Consistently, WASP deficient cells displayed lower levels of NPM1 and a heightened inflammatory cytokine response to LPS, which was rescued by overexpressing NPM1. Together, our research provides novel insights into the critical role of WASP in nucleolar function and the modulation of inflammatory cytokine production.

## INTRODUCTION

WAS is an X-linked recessive disease caused by mutations in the *WAS* gene that encodes WASP. The disease is characterized by a range of clinical features, including microthrombocytopenia, recurrent infections, immunodeficiency, autoimmunity, and malignancy ^1^. WASP is primarily expressed in hematopoietic cells and is an important nucleation-promoting factor that plays a critical role in regulating actin-polymerization ^2^. Although abnormalities in actin cytoskeleton-dependent cellular processes are well-established in the pathogenesis of WAS, studies indicate that both murine models and patients with enhanced actin polymerization due to WAS mutations still display classical features of the disease ^3,4^. This suggests that WASP has other roles beyond regulating actin nucleation. Studies from our group and others have demonstrated the WASP functions in multiple biological processes, including regulating message RNA (mRNA) alternative splicing ^5^, maintaining genome stability ^6,7,8^, nuclear translocation of NFAT2 and NF-κB (RelA) ^9^, and TH1 lineage commitment ^10^.

The emergence of induced pluripotent stem cell (iPSC) technology offers a promising solution to overcome the constraints in studying WAS. iPSCs can differentiate into various cell types relevant to WAS and provide an unlimited source of genetically well-defined experimental materials ^11^. Using previously characterized isogenic macrophage models generated from WASP knockout iPSCs, we investigated the nuclear function of WASP ^5^. The nucleolus is a non-membrane-bound organelle that consists of three compartments formed by liquid-liquid phase separation (LLPS), namely fibrillar center (FC), dense fibrillar component (DFC), and granular component (GC). Functionally, it plays a critical role in ribosomal biogenesis ^12^. Within the nucleolus, ribosomal DNA (rDNA) is transcribed by RNA polymerase I (PoI I) into the 47S pre-rRNA, which then undergoes various processing steps to yield mature rRNA molecules, including 5.8S, 18S, and 28S rRNA. These mature rRNA molecules are then assembled with ribosomal proteins and 5S rRNA to form fully functional mature ribosomes ^13,14^. Interestingly, studies have linked the nucleolus with viruses and bacterial infections, highlighting the immunity function of this organelle ^15,16^. The Las17, the yeast homolog of WASP, has been shown to play a role in maintaining nucleolar structure. The *Las17Δ* mutant results in various morphological changes in the nucleolus, indicating its importance in nucleolar structure maintenance ^17^. However, it remains unknown if WASP is present within the nucleolus and whether it can regulate nucleolar function and structure in humans.

Our findings revealed that a fraction of WASP is localized within the nucleolus and interacts with many nucleolar proteins, including NPM1 and FBL. The depletion of WASP in macrophages resulted in a significant reduction in nucleolar size and disrupted nucleolar morphology, along with decreased rRNA transcription. Moreover, the nucleolar and rRNA transcription phenotypes were confirmed in patient samples. Importantly, the loss of WASP was positively correlated with decreased NPM1 expression levels and augmented production of inflammatory cytokines in response to LPS, which can be rescued by overexpressing NPM1. These findings highlight a previously unknown role of WASP in regulating the nucleolar homeostasis and modulating the production of inflammatory cytokines, providing novel insights into the molecular mechanisms underlying WAS.

## RESULTS

### WASP is localized in the nucleolus

To investigate the subcellular localization of WASP within the nucleus, we performed ultrastructural analysis at the single-cell level. Consistent with previous studies ^5,18^, WASP signals were present inside the nucleus. Interestingly, our findings revealed clear nucleolar localization of WASP (Fig. 1A, and Supplementary Fig. S1A, B). A further immunofluorescence analysis of endogenous WASP demonstrated that a portion of WASP signals co-localized with nucleolar proteins NPM1 in the GC layer, FBL in the DFC, and RPA-194 in the FC ^19^ (Fig. 1B, C). These orthogonal results establish WASP as a newly identified nucleolar protein, suggesting functional roles of WASP in this multifaceted nuclear organelle.

**Fig. 1.**
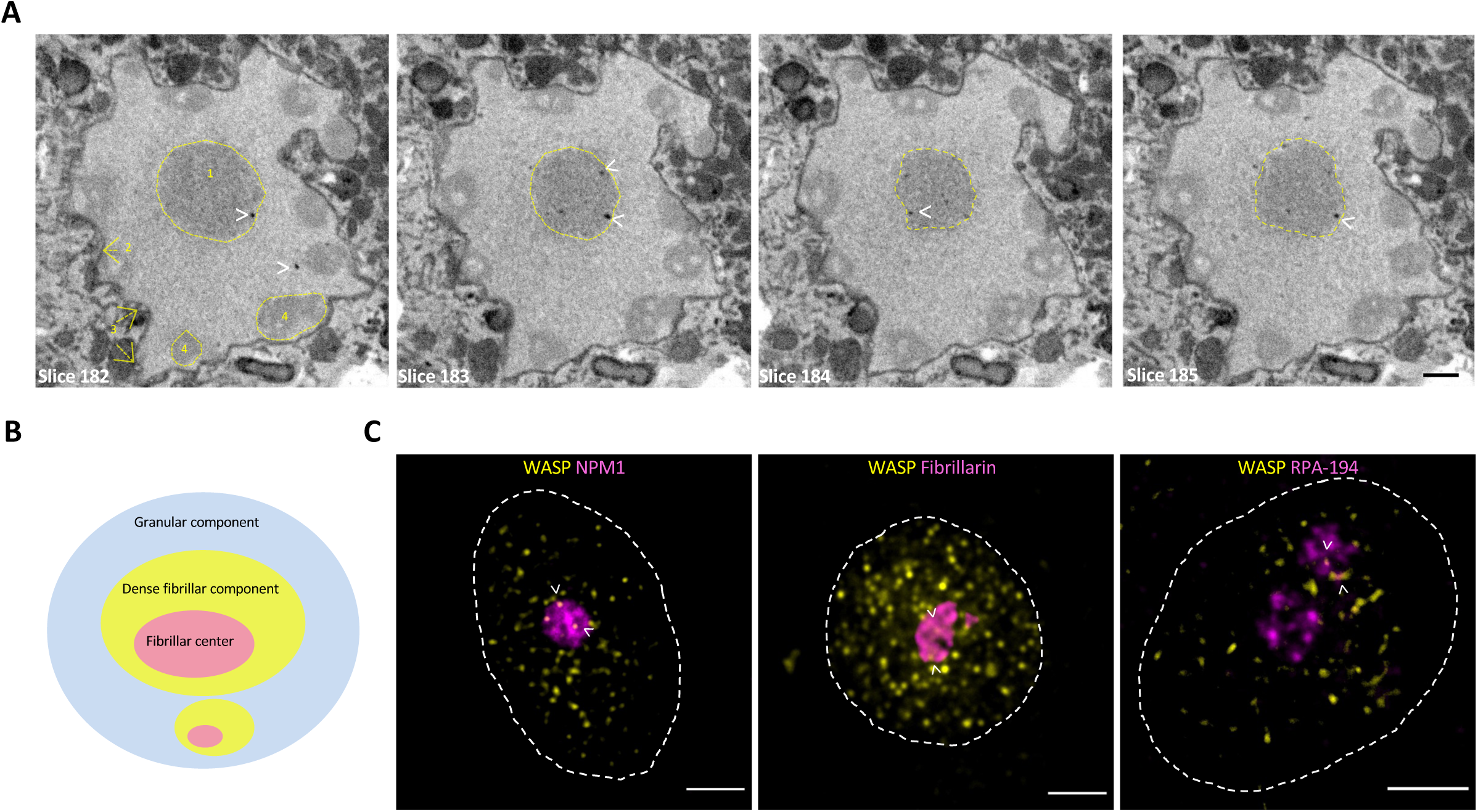
WASP is a nucleolar protein. A, SBF-SEM images of WT B cells expressing a WASP-miniSOG fusion protein. The sub-nuclear localization of WASP was visualized using the photo-oxidation of DAB by miniSOG (see Methods). High-contrast dark spots, indicated by white arrowheads, represent the WASP signals. Yellow arrows and outlines indicate nuclear landmarks: 1, nucleolus; 2, nuclear envelope; 3, nuclear pores; 4, heterochromatin. Bar = 1 μm. B, A schematic diagram of the mammalian nucleolus highlighting its three sub-domains: granular component (GC), dense fibrillar component (DFC), and fibrillar center (FC). C, Representative immunofluorescent images of iMPs showing WASP (yellow) and the subdomains of nucleoli: NPM1 for GC, Fibrillarin for DFC, and RPA-194 for FC. White arrowheads indicate the overlap between WASP and the sub-nucleolar structure. White dash line indicates the nuclear envelope. Bar = 3 μm.

### WASP interacts with NPM1 and FBL

To explore the nucleolar role of WASP further, we investigated if WASP could physically interact with nucleolar proteins. Using a proteomics approach, we identified 112 nucleolar proteins that interact with WASP ^5,20^ (Fig. 2A, Table 1). These potential WASP partners in the nucleolus are significantly enriched in gene ontology (GO) terms associated with biological processes involving the ribosome (Fig. 2B), which is consistent with the well-established role of the nucleolus in ribosomal biogenesis ^21^ and suggests that WASP is involved in the regulation of ribosomal function. Two essential nucleolar proteins, NPM1 and FBL, were identified as potential interaction partners of WASP. To confirm these interactions, we conducted a proximity ligation assay (PLA) experiment, which allows the detection of endogenous protein-protein interactions within a distance of less than 40 nm ^5^. PLA signals from WASP-NPM1 and WASP-FBL were found to be distributed within the nucleolus, providing strong evidence for a physical interaction between WASP and these known nucleolar proteins (Fig. 2C). Thus, we conclude that WASP localizes in the nucleolus and interacts with nucleolar constituent proteins therein.

**Fig. 2.**
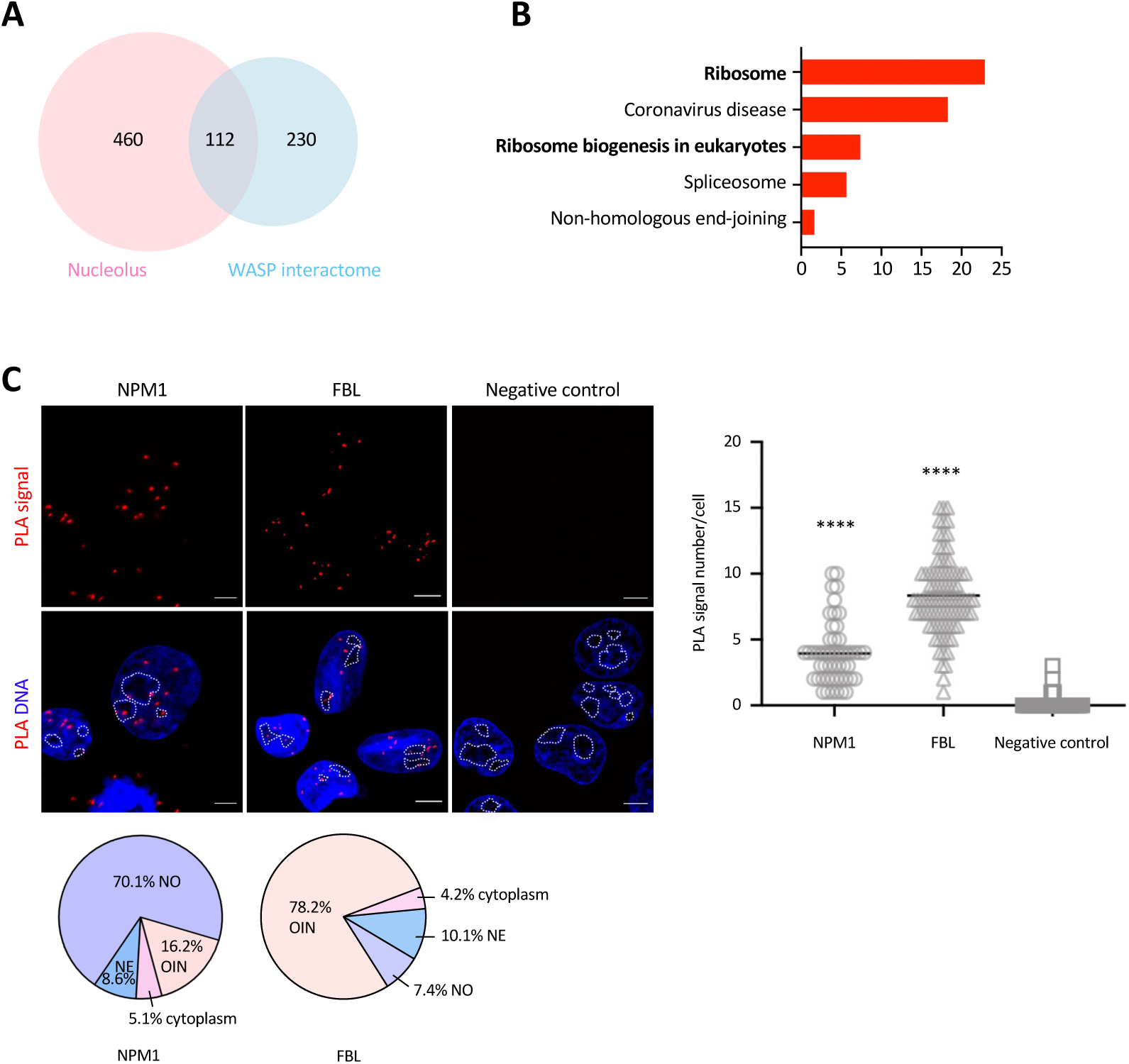
WASP binds nucleolus proteins. A, Venn diagram showing shared and distinct proteins identified in the interactome of WASP and the nucleolus. B, GO analysis of the overlap between WASP partners and nucleolar proteins according to reference ^20^. C, Representative immunofluorescence images of proximity ligation assay (PLA) signals for WASP-NPM1 and WASP-FBL in WT B cells. The pie chart illustrates the percentage of PLA signals detected in various sub-cellular locations. PLA signals are found in the expected sub-cellular locations, including the nuclear envelope (NE), nucleolus (NO), and other locations inside the nucleus (OIN). White dash line indicates the shape of the nucleolus. Right: quantitative analysis of PLA signals. n = 50 NPM1-WASP, n = 87 for FBL-WASP, and n = 49 for negative control. **** p < 0.0001. Nuclei were stained with DAPI. Bar = 5 μm.

**Table 1.**
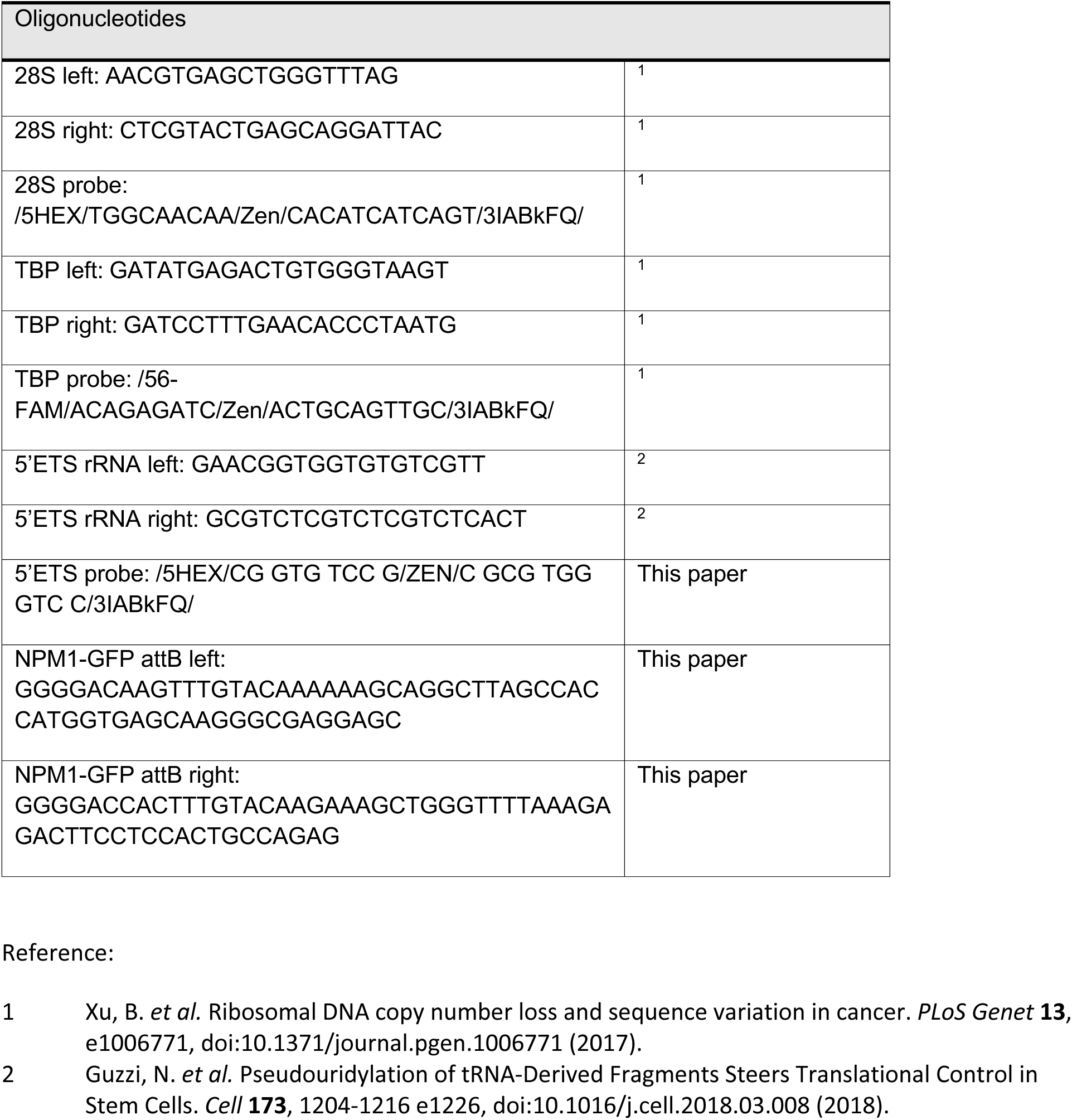

### WASP deficiency alters nucleolus architecture

To investigate the role of WASP in the nucleolus, we utilized wild-type (WT) iPSC-derived macrophages (iMPs) and isogenic WASP-KO iMPs ^5^ to examine whether WASP affects sub-nucleolar structure by super-resolution imaging techniques. We found that the localization of the markers of the three sub-nucleolar compartments were unaffected by the knockout, suggesting no gross disruption of the main sub-nucleolar compartments (Supplementary Fig. S2A). However, quantitative analysis revealed a significant reduction in nucleolar size, as indicated by the GC marker NPM1, in WASP-KO iMPs (Fig. 3A, B). Quantitative fluorescence imaging analysis also showed significantly lower levels of NPM1 expression in WASP-KO iMPs (Fig. 3A, B, E). Additionally, nucleoli in WASP-KO iMPs exhibited a loss of the smooth liquid-droplet-like appearance (Fig. 3A, B). These morphological changes were confirmed using another sub-nucleolar marker, FBL (Fig. 3C, D), whose expression level was similar between WT iMPs and WASP-KO iMPs (Supplementary Fig. S2B, C).

**Fig. 3.**
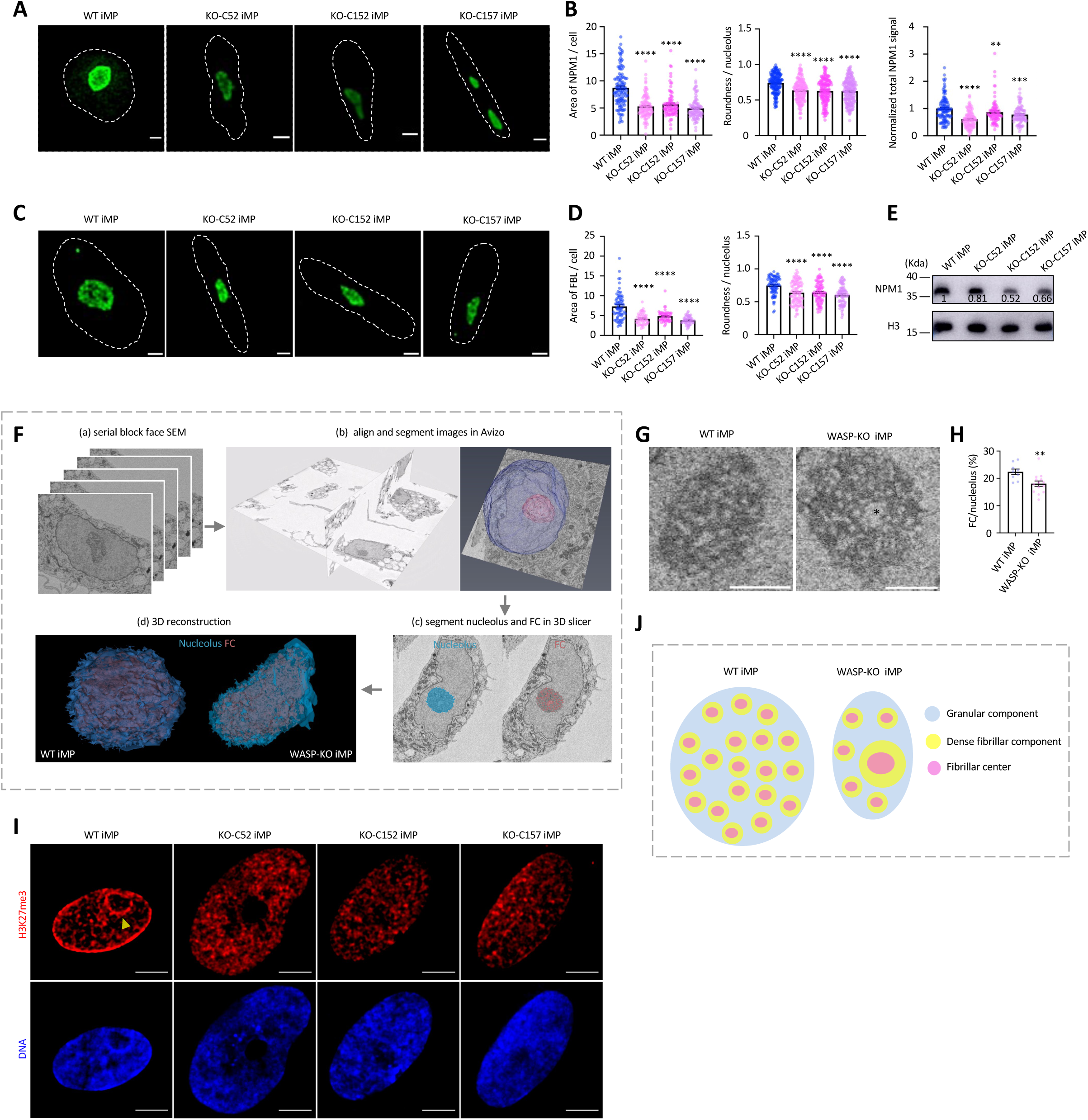
WASP deficiency affects nucleolar structure. A, Representative maximum projection confocal images of NPM1 in WT and three WASP KO macrophages. White dash lines indicate the shape of the nucleus. Bar = 2 μm. B, Quantitative analysis of the NPM1 signals in (A). The area of NPM1 per cell and normalized total NPM1 signals were analyzed: n = 112 (WT iMP), n = 106 (KO-C52 iMP), n = 87 (KO-C152 iMP), n = 96 (KO-C157 iMP). The roundness of NPM1 per nucleolus was also analyzed: n = 176 (WT iMP), n = 177 (KO-C52 iMP), n = 177 (KO-C152 iMP), n = 211 (KO-C157 iMP). Two biological replicates were performed independently. For the normalized NPM1 signals, WT signals were set as 1, and the WASP-KO signals were normalized to WT signals. Data are presented as mean ± SEM. Statistical analysis was performed using the Mann-Whitney test. **** p < 0.0001. C, Representative maximum projection confocal images of FBL in WT and three WASP KO macrophages. The white dash line indicates the shape of the nucleus. Bar = 2 μm. D, Quantitative analysis of FBL signals in (C). The area and roundness of FBL per cell were analyzed: n = 71 (WT iMP), n = 56 (KO-C52 iMP), n = 52 (KO-C152 iMP), n = 52 (KO-C157 iMP). Three biological replicates. Data are shown as mean ± SEM. Mann-Whitney test. **** p < 0.0001. E, Representative Western blot analysis of the NPM1 expression in WT iMP and three WASP-KO iMPs. H3 was used as a loading control. The normalized expression value of NPM1 relative to WT iMP is indicated by the numbers. The experiment was repeated twice independently. F, The schematic diagram illustrates the segmentation process of the FC within the nucleolus. (a) Image slices were aligned using Avizo software. (b) The FC and nucleolus were segmented using Avizo software. (c) A nucleolus segmentation algorithm was developed with 3D Slicer software, enabling the precise segmentation of the nucleolus and FC based on the previously obtained segmented nucleus in step (b). (d) 3D images of WT iMP and WASP-KO iMPs were generated using 3D Slicer software. G, Representative SBF-SEM images of nucleolus in WT iMP and WASP-KO iMPs. Bar = 1 μm. The asterisk indicates the presence of one large-sized FC. H, Quantitative analysis of the percentage of FC volume per nucleolus in WT iMP and WASP-KO iMP. n = 9 (WT iMPs), n = 14 (WASP-KO iMPs). Mann-Whitney test ** p = 0.0055. I. Confocal immunofluorescence images show the histone modification marker H3K27me3 in WT iMPs and KO iMPs (3 clones). Nuclei were stained with DAPI. Prominent perinuclear heterochromatin outside of the nucleolus is indicated by yellow arrowheads. Scale bar = 5 μm. J, Schematic diagram illustrating the differences in nucleolar structure of WT and WASP-KO iMPs.

We utilized serial block-face scanning electron microscopy (SBF-SEM) to examine the three-dimensional architecture of nucleoli in both WT and WASP-KO iMPs (Fig. 3F, Supplementary Movie). The nucleolus of both genotypes appeared as a reticulated structure (Fig. 3G). It has been reported that cells with low transcription activity often have small nucleoli containing one large FC ^22^. Notably, WASP-KO iMPs exhibited small nucleoli (Fig. 3A-D) and the presence of a single prominent FC in 44% (4 out of 9) of WASP-KO iMPs. This structural anomaly, which was absent in 100% (7 out of 7) of WT iMPs, suggests a decrease in transcription levels in WASP-KO iMPs. Additionally, our findings demonstrated a significant reduction in the ratio of the FC territory per nucleolus in WASP-KO iMPs (Fig. 3H). Lastly, perinucleolar heterochromatin, a shell of highly condensed heterochromatin around the nucleolus ^23^, was absent in WASP-KO iMPs (Fig. 3I).

Taken together, our results demonstrate that cells deficient in WASP exhibit smaller and misshapen nucleolus, along with the absence of perinucleolar heterochromatin and decreased NPM1 expression levels. Additionally, WASP-KO iMPs displayed a reduced proportion of FC and the presence of a single prominent FC (Fig. 3J). These findings strongly suggest that WASP is a nucleolar protein critical for maintaining nucleolar morphology and organization in macrophages.

### WASP deficiency impairs rRNA transcription and inflammatory cytokine production

The nucleolus is a critical compartment for ribosome biogenesis, where precursor 47S rRNA is synthesized by RNA Pol I and processed into mature rRNA. During rRNA maturation, the elimination of 5’ external transcribed spacer (5’ETS) is essential for the primary transcript step, allowing the maturation of 5.8S, 18S, and 28S rRNA ^14,24^. To explore the function of WASP in rRNA transcription, we first performed qRT-PCR to evaluate the level of different rRNA transcripts in WT and WASP-KO iMPs. We found that WASP knockout led to lower levels of the 5’ ETS and 28S rRNA (Fig. 4A). To further investigate the effect of WASP knockout on rRNA transcription in situ, we performed a 5-ethynyluridine (EU) pulse labeling experiment to quantify newly synthesized RNA (most of which is rRNA) and found significantly less nascent rRNA in WASP-KO iMPs (Fig. 4B), suggesting WASP knockout reduced the rate of rRNA transcription.

**Fig. 4.**
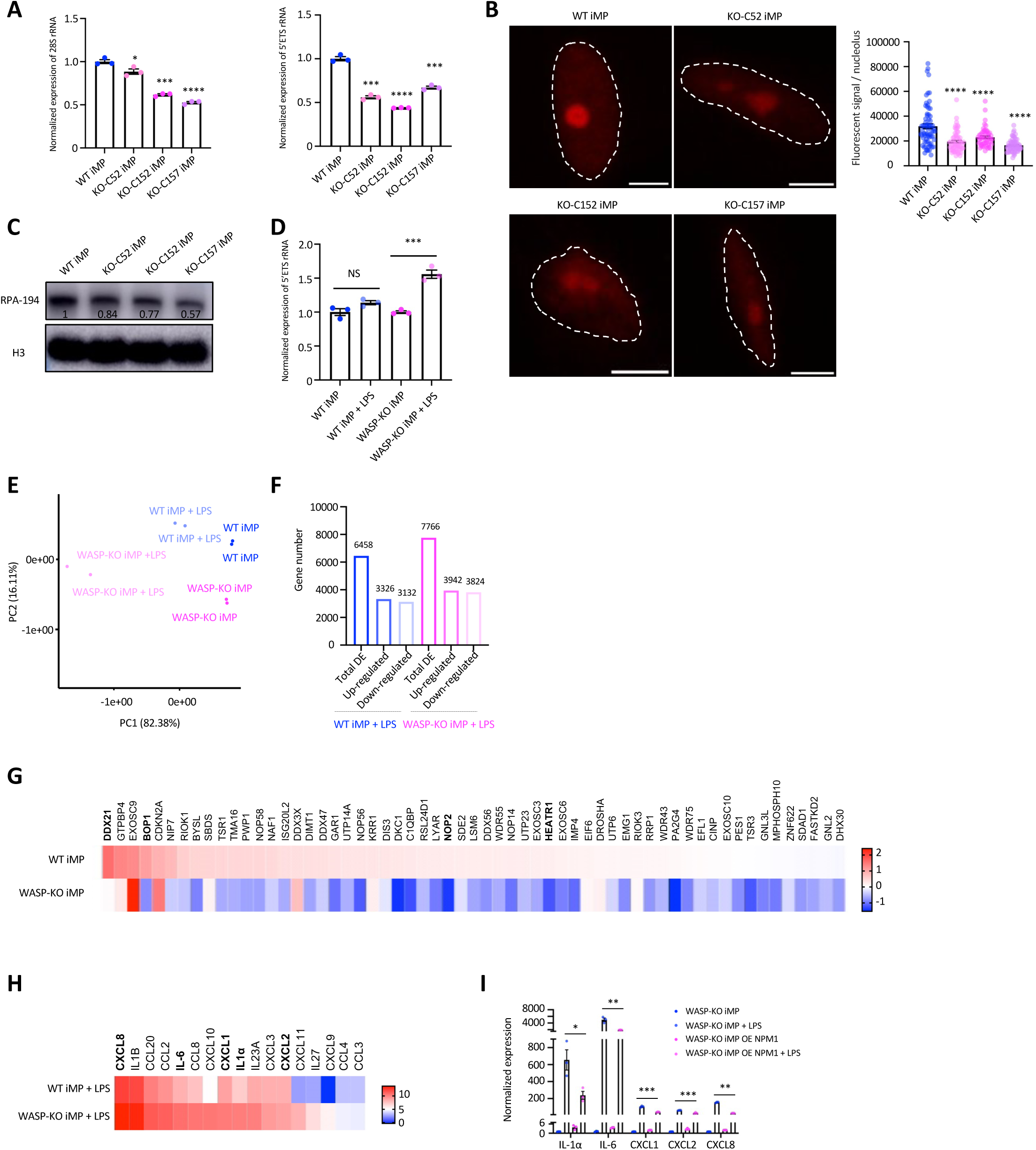
Impaired ribosomal RNA transcription in WASP-deficient cells. A, Representative qPCR analysis of 28S rRNA and 5’ETS, with more than three biological repeats. Data are presented as mean ± SEM and analyzed using Student’s t-test. For 28S rRNA, * p = 0.0383, *** p = 0.0001. For 5’ETS, *** p = 0.0001 for KO-C52 iMP, *** p = 0.0003 for KO-C157. **** p <0.0001 B, Representative images of EU-labeled nascent RNA. The total fluorescence signal per nucleolus was analyzed for n = 70 (WT iMPs), n = 60 (KO-C52 iMPs), n = 51 (KO-C152 iMPs), n = 66 (KO-C157 iMPs), with biological replicate n = 3. Data are presented as mean ± SEM and analyzed by Mann-Whitney test (**** p <0.0001). C, Representative Western blot analysis of RPA-194 expression in WT iMP and three WAS KO-iMPs, with H3 as a loading control. The numbers indicate the normalized expression value of RPA-194 relative to WT iMP, with more than three biological repeats. D, Normalized expression of 5’ETS upon the stimulation of iMPs with LPS for 6 hours, with more than three biological repeats. Data are presented as mean ± SEM and analyzed using Student’s t-test. *** p = 0.0009. E, PCA plot of RNA-seq samples. F, The number of genes showing differential expression after LPS stimulation. G, Heatmap illustrating the log2-fold change of expression levels of genes associated with ribosome biogenesis and rRNA processing in WT iMPs and WASP-KO iMPs following LPS stimulation, using the gene list from Supplementary Fig. S3D. Initial calculations involved obtaining the mean FPKM from two biological replicates. Subsequently, the fold change was computed by dividing the (FPKM + 1) after LPS stimulation by the value from the naive group. H, Heatmap illustrating the log2-fold change of genes associated with the inflammatory cytokine production after LPS stimulation. The values in the heatmap represent the mean FPKM from two biological replicates. I, Normalized qPCR analysis was performed to assess IL-1α, IL6, CXCL1, CXCL2, and CXCL8 in WASP-KO cells, with/without LPS stimulation and/or NPM1 overexpression (OE). Data are presented as mean ± SEM, and statistical significance was determined using Student’s t-test (n=3). Comparisons include WASP-KO iMPs with LPS vs. WASP-KO OE NPM1 iMPs with LPS. Significance levels are indicated as follows: IL-1α * P = 0.0312, IL6 ** P = 0.004, CXCL1 *** P = 0.0005, CXCL2 ** P = 0.004, and CXCL8 **** P < 0.0001.

To rule out the possibility that the reduced level of rRNA is due to a lower copy number of rDNA, we conducted droplet digital PCR (ddPCR) experiments and found no significant difference in rDNA copy number between WT and WASP-KO iMPs, suggesting that WASP regulates rRNA transcription (Supplementary Fig. S3A). Western blots showed lower RNA Pol I expression in WASP-KO iMPs (Fig. 4C), which is consistent with the reduction of the relative FC territory in the nucleolus (Fig. 3H) and lower transcriptional levels of rRNA (Fig. 4A, B).

WAS patients suffer from recurrent infections, attributable in large part to the defects in innate immune cells such as macrophages in sensing bacterial infections and/or clearing exogenous pathogens ^25^. Recent studies have revealed that nucleolar proteins regulate innate immune response ^15,16^, suggesting the nucleolus as a cellular organelle involved in innate immunity. Thus, we decided to explore the functional connections between the newly discovered nucleolar role of WASP and WAS innate immunodeficiency using WT and WASP-KO iMPs challenged by LPS. LPS stimulation led to an increased level of TNF-α secretion, confirming the validity of the treatment regime (Supplementary Fig. S3B). Interestingly, WT iMPs showed no changes in rRNA levels 6 hours after LPS treatment and reduced rRNA levels at 12 hours, whereas WASP-KO iMPs showed significantly elevated rRNA levels, which increased approximately 50% 6 hours after LPS treatment and remained high at 12 hours (Fig. 4D, and Supplementary Fig. S3C). To further investigate the full spectrum of gene expression changes associated with this phenotype, we performed deep RNA-seq of two sets of biological replicate samples. Biological replicates of all experimental groups clustered tightly together in principal component analysis (PCA), indicating good reproducibility. Most variance in PC1 (82.38%) was explained by the difference between naïve and LPS treated WASP-KO iMPs, which dwarfs the difference between naïve and LPS treated WT iMPs and that between naïve WASP-KO iMPs and WT iMPs (Fig. 4E). This finding suggests the involvement of WASP in shaping macrophage response to LPS. LPS stimulation resulted in 6458 differentially expressed genes (DEGs) in WT iMPs, out of which 3326 genes were upregulated (FDR<0.05). In WASP-KO iMPs, 3942 out of 7766 DEGs showed upregulation upon LPS stimulation (Fig. 4F). Intriguingly, GO analysis showed only WT iMPs upregulated genes enriched in ribosome biogenesis and rRNA processing after LPS stimulation (Supplementary Fig. S3D, E). Furthermore, detailed analysis showed that 83.1% of genes upregulated in the rRNA processing pathway and ribosome biogenesis pathway in WT iMP cells were downregulated in WASP-KO iMPs (Fig. 4G). For example, well-known genes involved in rRNA processing and ribosome biogenesis, such as *DDX21* ^26^, *BOP1* ^27^, *NOP2* ^28^ and *HEATR1* ^29^, are only up-regulated in the WT iMPs (Fig. 4G). Collectively, our findings highlight the critical role of WASP in regulating rRNA transcription and/or processing.

The upregulated DEGs in LPS-stimulated WT and WASP-KO iMPs were unsurprisingly enriched in biological processes associated with innate immunity and inflammatory response (Supplementary Fig. S3D, E). Since we found lower levels of NPM1 in WASP deficient cells (Fig. 3A-B, E) and reduced NPM1 expression is known to be associated with elevated inflammatory cytokine release following LPS stimulation^30^, we therefore examined inflammatory cytokine production upon LPS stimulation. Interestingly, WASP-deficient cells exhibited elevated levels of inflammatory cytokines (Fig. 4H). We hypothesized that overexpressing NPM1 could alleviate this heightened cytokine response. To test this hypothesis, we established an inducible system for NPM1 expression and verified that the overexpressed NPM1 had the same subcellular localization as endogenous NPM1 (Fig. S3F). Remarkably, overexpression of NPM1 significantly decreased LPS-stimulated production of all tested cytokines in WASP-KO iMPs (Fig. 4I). Together, these results support the notion that the nucleolar function of WASP underlies normal innate immune response in macrophages.

### Confirmation of nucleolar and rRNA phenotypes in patient samples

To validate the findings in iMP models, we examined nucleolar morphology, protein expression, and rRNA transcription in peripheral blood of a WAS patient and his first-degree relatives. The patient presented with symptoms including thrombocytopenia, intermittent diarrhea, and a skin rash resembling eczema. Confirmatory diagnosis of WAS was established through whole exome sequencing, which revealed a point mutation (NM_000377.3: exon4: c.397G>A Glu133Lys) resulting in a missense mutation. Western blot (Fig. 5A) of PBMCs confirmed the absence of WASP expression in the patient. Consistent with our findings in WT and WASP-KO iMPs, macrophage cells derived from patient PBMCs exhibited significantly smaller nucleoli, decreased roundness, and lower total NPM1 signals (Fig. 5B, C). Moreover, we observed a substantial reduction in EU labeled nascent rRNA in patient macrophage cells (Fig. 5D, E). These findings show that the nucleolar changes and altered rRNA transcription observed in isogenic iMP models are bona fide WAS pathological features.

**Fig. 5.**
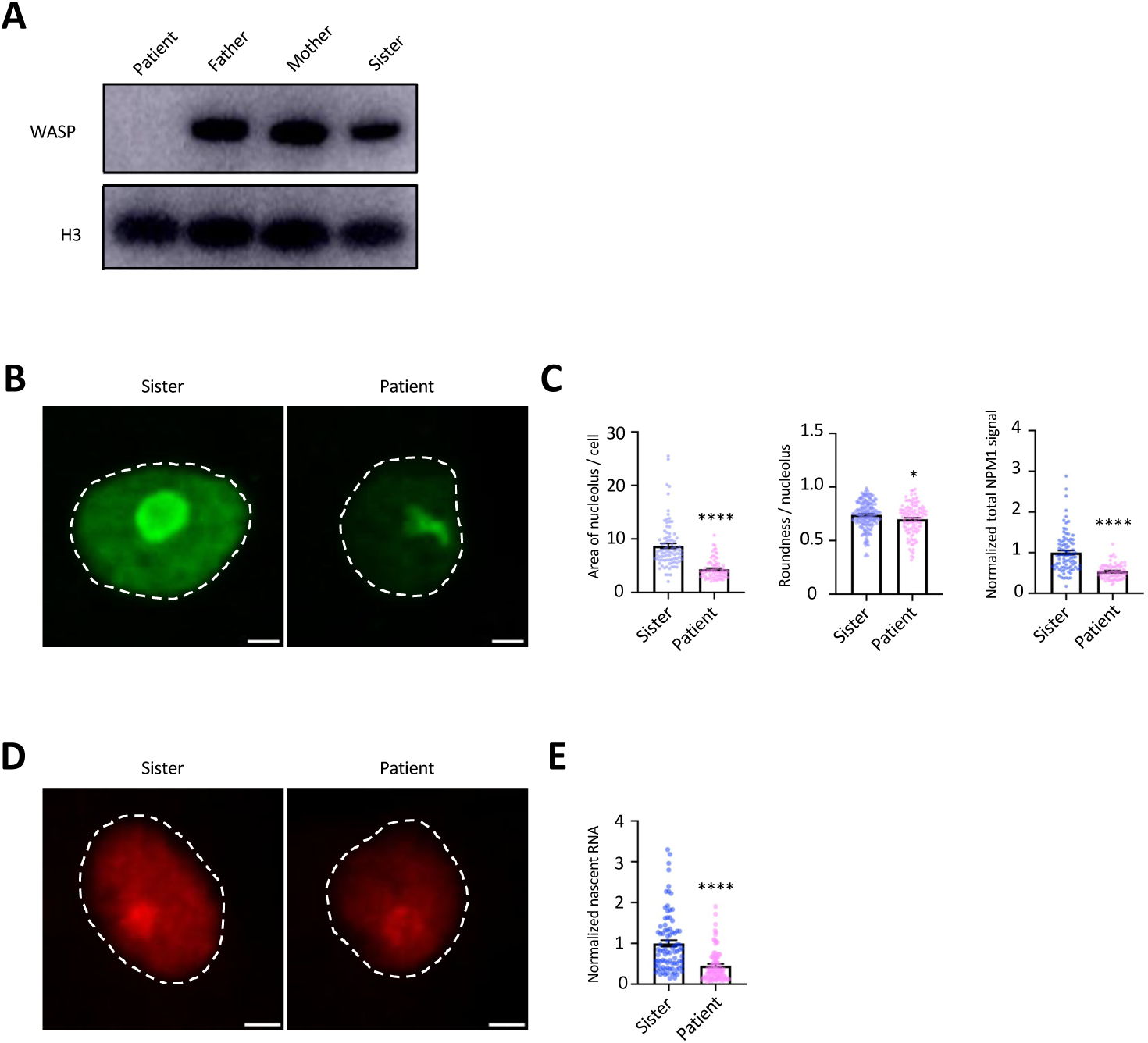
Recapitulation of nucleolar structure changes and ribosomal RNA profile in patient samples. A, Western blot of WASP expression levels in PBMC samples from the WAS patient, his parents, and sister. H3 was used as a loading control. B, Representative maximum projection confocal images of the NPM1 in sister and WAS patient PBMC-derived macrophages are shown. The white dashed line indicates the shape of the nucleus. Bar = 2 μm. C, The area of NPM1 per cell and the normalized total NPM1 signals were analyzed, with a sample size of n = 88 for sister and n = 80 for patient. The roundness of NPM1 per nucleolus was also analyzed, with a sample size of n = 197 for sister and n = 114 for patient. The signal from the sister was set as the man value as 1, and the signal from the patient was normalized to the signal from the sister. Data are presented as mean ± SEM. Statistical analysis was performed using the Mann-Whitney test. **** p < 0.0001, * p = 0.0278. D, Representative maximum projection confocal images of newly synthesized RNA in sister and WAS patient PBMC-derived macrophages. The white dashed line indicates the shape of the nucleus. Bar = 2 μm. E, Representative confocal images of EU-labeled nascent RNA. The total fluorescence signal per nucleus was normalized to macrophage cells from sister and analyzed for a sample size of n = 86 for sister and n = 82 for patients. Data are presented as mean ± SEM and were analyzed by the Mann-Whitney test (**** p < 0.0001).

## DISCUSSION

Increasing evidence suggests that WASP has many nuclear functions, but its nucleolar involvement remains unexplored. In this study, we utilized a previously established isogenic WASP-KO iMP model to investigate the potential role of WASP in the nucleolus. By employing SBF-SEM and immunofluorescence, we provide the first evidence that WASP is a nucleolar protein (Fig. 1 and supplementary Fig. S1). Additionally, we identified 112 nucleolar proteins as WASP interaction partners (Fig. 2A) and further validated the interactions between WASP and NPM1 and FBL (Fig. 2C). These findings suggest that the nucleolar localization of WASP is conserved across yeast and mammalian cells ^17^.

An intriguing observation in our study was the reduced size and irregular shape of the nucleolus in WASP-deficient cells (Fig. 3A-D, Fig. 5B, C). The presence of small nucleoli accompanied by one large-sized FC has been associated with lower transcription levels ^22^. The same morphology was displayed by WASP-KO cells (Fig. 3A-D, G, H) and accompanied by lower rRNA transcription levels (Fig. 4A, B). These findings suggest a link between the observed structural alterations in nucleoli and the reduced transcriptional activity of rRNA in WASP-KO cells. A decrease in nucleolar size is associated with active bacterial infection and is suggested to be a specific response to bacterial challenge ^15^. Additionally, exposure to heat-shock stress has been linked to the loss of nucleolar roundness ^19^. Previous studies have demonstrated that NPM1-knockdown cells exhibit smaller and irregular nucleoli ^31^. We observed similar structural changes and lower NPM1 expression levels in WASP-KO cells (Fig. 3 A-E, Fig. 5B, C). Intriguingly, we previously showed that WASP deficiency causes abnormal morphology of the nuclear speckle, another non-membrane bound organelle formed by LLPS, where wild-type WASP interacts with the nuclear speckle protein SRSF2 and constrains its mobility, providing a possible mechanism for the phenotype ^5^. Drawing an interesting parallel, this study shows WASP interacts with nucleolar proteins NPM1 and FBL. It is of interest to study the functional significance of these interactions in nucleolus organization and rRNA transcription in the future. Taken together, the observed structural changes in WASP-KO iMPs suggest a role for WASP in nucleolar dynamics and underscore its importance in maintaining the nucleolar structure.

In addition, we evaluated the impact of WASP on rRNA production. We observed that WASP-deficient cells exhibited lower rRNA production (Fig. 4A, B, Fig. 5D, E) and decreased RNA Pol I expression (Fig. 4C). One possible mechanism by which WASP regulates rRNA transcription is through its interaction with NPM1, a known regulator of pre-rRNA cleavage and rRNA production^32,33^.

Translation inhibition has been recognized as an effective antiviral immune response, as it can block viral gene expression following viral infections ^33,34^. Similarly, bacterial infection effectors have been shown to induce host translation arrest, leading to a stronger immune response ^15,35,36^. In the present study, we observed that WT iMPs tightly regulated rRNA expression levels after LPS stimulation, while WASP-KO iMPs failed to do so and exhibited significantly higher levels of rRNA after LPS challenge (Fig. 4D). Additionally, WASP-KO iMPs displayed abnormal regulation of genes involved in ribosome biogenesis and rRNA processing upon LPS stimulation (Fig 4G and Supplementary Fig. S3D, E). These findings suggest that WASP regulates rRNA processing and provide a plausible explanation of the elevated levels of rRNA in WASP-deficient cells.

Studies have highlighted the crucial role of the nucleolus in maintaining immune homeostasis, particularly in response to infections ^15,16^. Tightly regulated expression of inflammatory cytokines is critical to proper immune response, but their excessive expression can lead to autoimmune diseases ^37,38^. NPM1 has been identified as a transcriptional co-repressor that negatively regulates cytokine production in LPS-stimulated macrophages ^30^. In line with these findings, WASP-KO iMPs that have lower NPM1 levels exhibited exacerbated inflammatory cytokine responses (Fig. 3H), which could be rescued by overexpression of NPM1 in the WASP-KO iMPs (Fig. 4I). The potential involvement of the nucleolus function of WASP in modulating inflammatory cytokine production provides a plausible explanation for the observed autoimmunity defects in WAS.

More importantly, all alterations in nucleolus and rRNA levels identified in WASP-KO iMPs are confirmed in patient samples (Fig. 5). Thus, the nucleolar defects of WASP deficient macrophages are hitherto unappreciated pathological features of WAS. The consistent findings in patient-derived macrophage cells support the notion that WASP plays a crucial role in regulating nucleolus architecture and ribosomal RNA transcription in a physiological context.

In conclusion, our study reveals that WASP resides in the nucleolus and regulates the structural integrity and rRNA metabolic processes in this organelle. The dysregulation of nucleolar functions in WAS-deficient cells suggests a potential link between the nucleolar dysfunction and immune-related defects observed in WAS. Further studies elucidating the underlying molecular mechanisms and exploring the broader impact of WASP on nucleolar dynamics and immune homeostasis in macrophages and other immune cell types will contribute to a better understanding of the pathophysiology of WAS and may uncover potential therapeutic targets for WAS and other immune-related disorders.

## METHODS

### Cell Culture

B-lymphocyte GM11518 cells were obtained from the Coriell Institute for Medical Research. The cells were cultured in Roswell Park Memorial Institute Medium 1640 supplemented with 15% fetal bovine serum.

iMP differentiation was carried out following a previously described protocol ^5^. Initially, iPSCs were cultured on MEF feeder for one to two generations. Then, iPSC clones were picked and seeded on OP9 feeders to initiate hematopoietic differentiation. After two weeks, differentiated iPSC cells were harvested. To generate macrophage cells, iPSCs were cultured with GM-CSF (Thermo Fisher, PHC2013) and M-CSF (Thermo Fisher, PHC9501) for two weeks and further matured with M-CSF for another week.

The study of human samples was approved by the Institutional Review Board (IRB) of the King Faisal Specialist Hospital & Research Center and the KAUST Institutional Biosafety and Bioethics Committee (IBEC). All of the human subjects signed the inform consent voluntarily after being clearly informed of all the content and details of the experiments. The patient is a 17-month-old boy who is experiencing symptoms including thrombocytopenia, intermittent diarrhea, and a skin rash resembling eczema. He was given a diagnosis with a WAS clinical score of 2 ^39^. Whole exome sequencing has confirmed the presence of mutations in the *WAS* gene. Specifically, the patient carries a point mutation (NM_000377.3: exon4: c.397G>A Glu133Lys) where one nucleotide (G) is substituted with another (A) on exon 4, resulting in a missense mutation from Glu to Lys. Peripheral blood mononuclear cells (PBMCs) were isolated from the patient’s blood and cultured in RPMI 1640 with 10% FBS (Gibco, 6140-079) for 24 hours to allow monocyte cells to attach. The non-adherent cells were then removed by washing with PBS. The remaining cells were cultured in RPMI 1640 with 10% FBS and 50 ng/mL M-CSF for one week, with medium changes every 3 days.

### Immunofluorescence Analysis

Cells were washed with PBS and fixed with 4% formaldehyde for 30 minutes. Next, they were permeabilized with 0.4% Triton X-100 for 15 minutes, followed by blocking with 6% normal donkey serum containing 0.1% Triton in PBS. The cells were then incubated with primary antibodies overnight at 4°C and subsequently with secondary antibodies for 45 minutes at room temperature. Finally, ProLong™ Glass Antifade Mountant (Thermo Fisher, P36980) was applied to the cells, which were then left to settle at room temperature overnight before imaging. Images were obtained using a Leica SP8 TCS STED 3X microscope with lightning.

### Serial Block-Face Scanning Electron Microscopy

MiniSOG^40^ (generously provided by R.Y. Tsien) was fused to the carboxyl terminus of WASP in a lentivirus expression vector. As a negative control, we used WASP fused to mEOS3, a fluorescence tag incapable of photo-oxidation of diaminobenzidine (DAB). Macrophages derived from normal cord blood adhered to 35 mm glass bottom grid-500 dishes (Ibidi, 81168) were fixed in 1% gluteraldehyde:1% paraformaldehyde in 0.15 M sodium cacodylate buffer, pH 7.4, containing 2 mM CaCl_2_ for 30 minutes at room temperature. The miniSOG mediated photo-oxidation of DAB was done as previously described ^41^. Samples were prepared for SBF-SEM as described in “NCMIR methods for 3D EM” (http://www.gatan.com/products/sem-imaging-spectroscopy/3view-system#resources). Briefly, they were then stained with 2% OsO_4_, 1.5% potassium ferrocyanide in 0.15 M cacodylate buffer, pH 7.4 containing 2 mM CaCl_2_ for 1 hour at 4°C. Samples were washed with deionized water (ddH_2_O), treated with thiocarbohydrazide for 20 minutes at room temperature, washed with ddH_2_O and stained again in 2% OsO_4_ for 1 hour at room temperature. Cell samples were counterstained with 1% uranyl acetate overnight at 4°C and subsequently washed with ddH_2_O before incubation in *en bloc* Walton’s lead nitrate at 60 degrees for 30 minutes. Samples were serially dehydrated with cold ethanol (35%, 50%, 70%, 90%, 100%, 100%) for 5 minutes for each step at 4°C, followed by cold absolute ethanol for 10 minutes on ice, and an additional incubation in absolute ethanol for 10 minutes at room temperature. Samples were serially infiltrated into Durcupan resin (11.4g part A, 10g part B, 0.3g part C, 0.075g part D) over the course of 6 hours (25:75 resin:ethanol, 50:50 resin:ethanol, 75:25 resin:ethanol) and left in 100% resin overnight. The next day, samples were incubated again with fresh Durcupan resin for 2 hours. To reduce potential sample charging while imaging, an additional half-batch of Durcupan resin (5.7g part A, 5g part B, 0.15g part C, 0.0375g part D) was made that contained 10mg of Polyanaline emeraldine base powder (Alfa Aesar, 32038). The samples were cured in the aforementioned resin at 60°C for 48 hours. Glass from the 35mm dish was etched off with hydrofluoric acid (Sigma Aldrich, 30107). The resin embedded ROIs that were marked during the photo-oxidation steps were excised using a jeweler’s saw, and then mounted (Electron Microscopy Sciences, 12642-14) onto a specimen pin (Gatan, 3VMRT12) using two-part conductive silver epoxy. The ROI was trimmed to yield a block face with dimensions ∼0.5 x 0.5mm and a thin layer of platinum was evaporated onto the block face using the Leica EM SCD 500 prior to imaging. Serial block face scanning electron microscopy was performed on a Zeiss Sigma VP (Cambridge, UK) equipped with a Gatan 3View XP2 (Pleasanton, CA, USA). The SEM was operated at 2.5 KeV with a 30 µm aperture in variable pressure mode set to 10 Pa pressure with a slice thickness of 50-70 nm. Images were taken at ∼10 nm/pixel with either a 6 µs dwell time using the 3View’s integrated backscatter detector with image dimensions ranging between 2848 x 2848 pixels and 3538 x 3538 pixels. Sectioning and imaging continued until the entire cell(s) of interest was imaged.

### Segmentation and 3D reconstruction of nucleolus

To segment the nucleus, the slices of the same sample were aligned using the Avizo software (2021.1) (Supplementary Movie). The sorted TIFF format images were then processed by OpenCV^42^ and Python 3.6. Images were denoised with the fast non-local means denoising algorithm (fastNlMeansDenoising) ^43^. The resized and denoised images were arranged into a 3D cube with dimensions of 640 x 640 x 640 and zero-padded. The cube was then converted into 640 2D images, each with dimensions of 640 x 640, along three orthogonal planes: x-y, x-z, and y-z. Using the raw images and their corresponding nucleus segmentation masks, we manually created three datasets, one for each plane (640 samples), to be used in the subsequent model training. For each plane, we fine-tuned a pre-trained U-Net provided by Tensorflow ^44^. The learning rate was set to 1e-4, the batch size was set to 1, and the number of epochs was set to 15.

To generate a consistent mask for the nucleus, we fine-tuned a pre-trained U-Net model to perform segmentation^45^. We analyzed the predicted masks along the x-y, x-z, and y-z planes. A pixel was considered part of the nucleus if it was predicted to be 1 in two or more planes. This process ensured consistency across the three planes and produced a more accurate segmentation of the nucleus. All model trainings and predictions were conducted on a workstation with 252 GB of RAM and 2 Tesla V100 GPUs.

Using the consistent mask generated in the previous step, we extracted the 3D nucleus from the original images for all samples. All 3D nuclei were further normalized with histogram equalization, which is a technique used to improve the contrast and brightness of an image by spreading out the intensity levels of the image histogram. We manually identified the approximate intervals of the nucleolus in the x, y, and z directions within the 3D cell nucleus volume. Next, we extracted the nucleolus region, applied a pixel-level cutoff of 100 to get the primary segmentation mask, and used erosion and dilation from scikit-image to remove noise^46^. This algorithm enabled us to accurately segment the fibrillar center structure of the nucleolus from the surrounding regions, thus obtaining a detailed representation of its shape and spatial distribution. Subsequently, we generated a mask of the nucleolus by filling the cavity structure based on the FC mask. With the segmentation masks of the nucleus, nucleolus, and the FC structure, we performed statistical calculations and visualized the 3D structure using 3D Slicer ^45^.

### Western blotting

To detect the protein expressions, we lysed cells in Laemmli sample buffer (Bio-Rad, 1610737) and loaded them onto 4-12% Bis-Tris Plus Gels (Thermo Fisher Scientific, NW04125BOX) for separation. Proteins were transferred to 0.2 µm polyvinylidene difluoride and blocked with 5% BSA. The antibodies and their concentrations used for the western blotting are listed in the table. We calculated the normalized expression value using the Gels model in Fiji (version: 2.3.0/1.53q).

### qPCR and Droplet Digital PCR (ddPCR)

RNA was extracted from the same amount of WT and WASP-KO iMPs using Trizol reagent (Life Technologies, 15596018). RNA samples were treated with DNase I (New England Biolabs, M0303S) to remove any remaining DNA. Then, the same volume of RNA was converted into cDNA to perform qPCR experiment. SsoAdvaned Universal SYBR Green Supermix (Bio-Rad, 1725270) was used for regular qPCR. GAPDH served as the control.

ddPCR was carried out using a QX200 Droplet Digital PCR system (Bio-Rad). For the rDNA copy number analysis, genomic DNA was isolated using a DNeasy Blood Tissue Kit (Qiagen, 69506). dd PCR was performed to measure 45S rDNA copy number as previously described^47^. 1 ng genomic DNA was digested with HaeIII (New England Biolabs, R0108L). TBP served as the internal control. The primer sequences are given in the Table 2.

### Nascent RNA labeling

To evaluate newly synthesized RNA, macrophages were labeled with 5-EU for 1 hour and detected using the Click-iT™ RNA Alexa Fluor™ 594 Imaging Kit (Invitrogen™ C10330). The images were captured using the Leica SP8 TCS STED 3X microscope with lightning.

### RNA-seq

RNA was isolated using Direct-zol RNA Miniprep Plus Kits (R2073). The quality of the RNA samples was evaluated using the Bioanalyzer (Agilent), with an RNA Integrity Number (RIN) above 9. The NEBNext® UltraTM Directional RNA Library Prep Kit for Illumina® (NEB, USA) was used to generate the sequencing library. RNA-seq was performed using the Illumina 6000 PE150 platform to generate paired-end 150 bp reads. The libraries were sequenced, resulting in approximately 12 GB of raw data reads per library.

### Bioinformatics analysis

The raw RNA-seq data were uploaded to the online RNA-seq data analysis website A.I.R. (https://transcriptomics.sequentiabiotech.com/) for further analysis. The sequencing reads were aligned to the hg38 human genome using the STAR alignment software (v2.5.1). To identify differentially expressed (DE) genes, we employed DESeq2, with a q-value threshold of <0.05. A Gene Ontology (GO) analysis was conducted using DAVID (v6.8) to provide a functional annotation and enrichment analysis of the DE genes.

### NPM1 overexpression and inflammatory cytokine production

To establish inducible NPM1 expression, the GFP-NPM WT plasmid was procured from Addgene (Plasmid #17578). The NPM1-GFP cDNA was cloned into pInducer20 (Plasmid #44012) utilizing the Gateway Cloning technology from Thermo Fisher, with primer details provided in the primer list.

A total of 500 ng of pInducer20-NPM1-GFP plasmid was mixed with Buffer R (derived from the Neon system kit, Cat#MPK1096) to create a final volume of 10 µl. Subsequently, 300,000 WASP-KO iMPs underwent electroporation using the Neon system with conditions set at 1600 V, 20 ms width, and 2 pulses. Following electroporation, the cells were promptly placed into separate wells of 6-well plates. After 24 hours of incubation, doxycycline (Thermo Fisher, Cat#BP2653-1) was introduced into the culture medium at a final concentration of 2 µg/ml to induce NPM1-GFP expression.

For subsequent experiments, a stimulation regimen was adopted: both WAS-KO iMPs and WAS-KO iMPs overexpressing NPM1-GFP were exposed to 1 µg/ml LPS for a 6-hour duration. Following the LPS stimulation period, RNA was extracted using Trizol reagent (Life Technologies, Cat#15596018) and subsequently reverse transcribed into cDNA using the iScript Reverse Transcription Supermix (BioRad, Cat#1708840). Quantitative PCR (qPCR) was performed using the CFX384 real-time PCR detection system (BioRad), utilizing TaqMan™ Fast Advanced Master Mix (Thermo Fisher, Cat# 4444557) and specific probes, including PGK1-VIC, IL1 α-FAM, IL6-FAM, CXCL1-FAM, CXCL2-FAM, and CXCL8-FAM, all obtained from Thermo Fisher.

### Statistical analysis

Results are reported as the mean ± standard error of the mean (SEM). Statistical comparisons were conducted using GraphPad Prism software and either the Student’s t-test or Mann-Whitney test. A p-value less than 0.05 was considered statistically significant.

### Data availability

The raw data and TMM value of the RNA-seq generated in this study were deposited in the NCBI GEO database.

## ACKNOWLEDGMENTS

We thank members of Li laboratory for valuable discussion and D. Chen for her administrative support. We thank Ali Reza Behzad of the KAUST Imaging and Characterization Core Lab, Thomas Theussl of the KAUST Visual Computing Center, and staff of Bioscience Core Lab for technical support. We thank Peter Karagiannis from KAUST for suggestions in writing. We also thank Dr. Corrado Calì for suggestions in image processing.

## Supplementary Figures

**Fig. S1.**
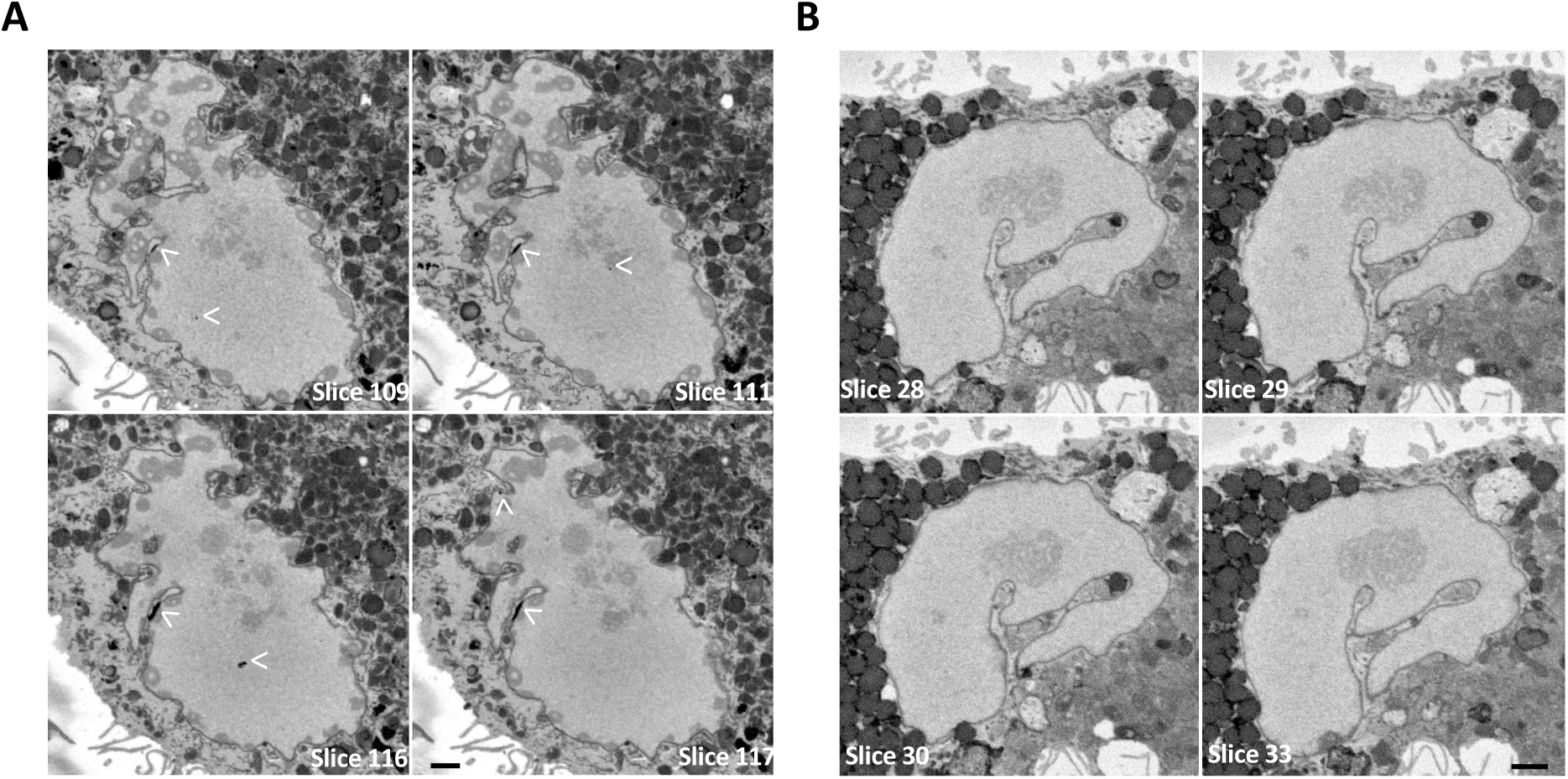
Nuclear localization of WT WASP, related to Figure 1. A, Sub-nuclear localization of WASP was visualized by the photo-oxidation of DAB by miniSOG. White arrowheads indicate WASP signals (high-contrast dark spots) at the nuclear envelope, nuclear periphery, and nuclear interior regions. B, SBF-SEM images of WT macrophages expressing a WASP-mEOS fusion protein, which is incapable of photo-oxidation of DAB and serves as a negative control for the specificity of the positive signals in A and Fig. S1A. Bar = 1 μm.

**Fig. S2:**
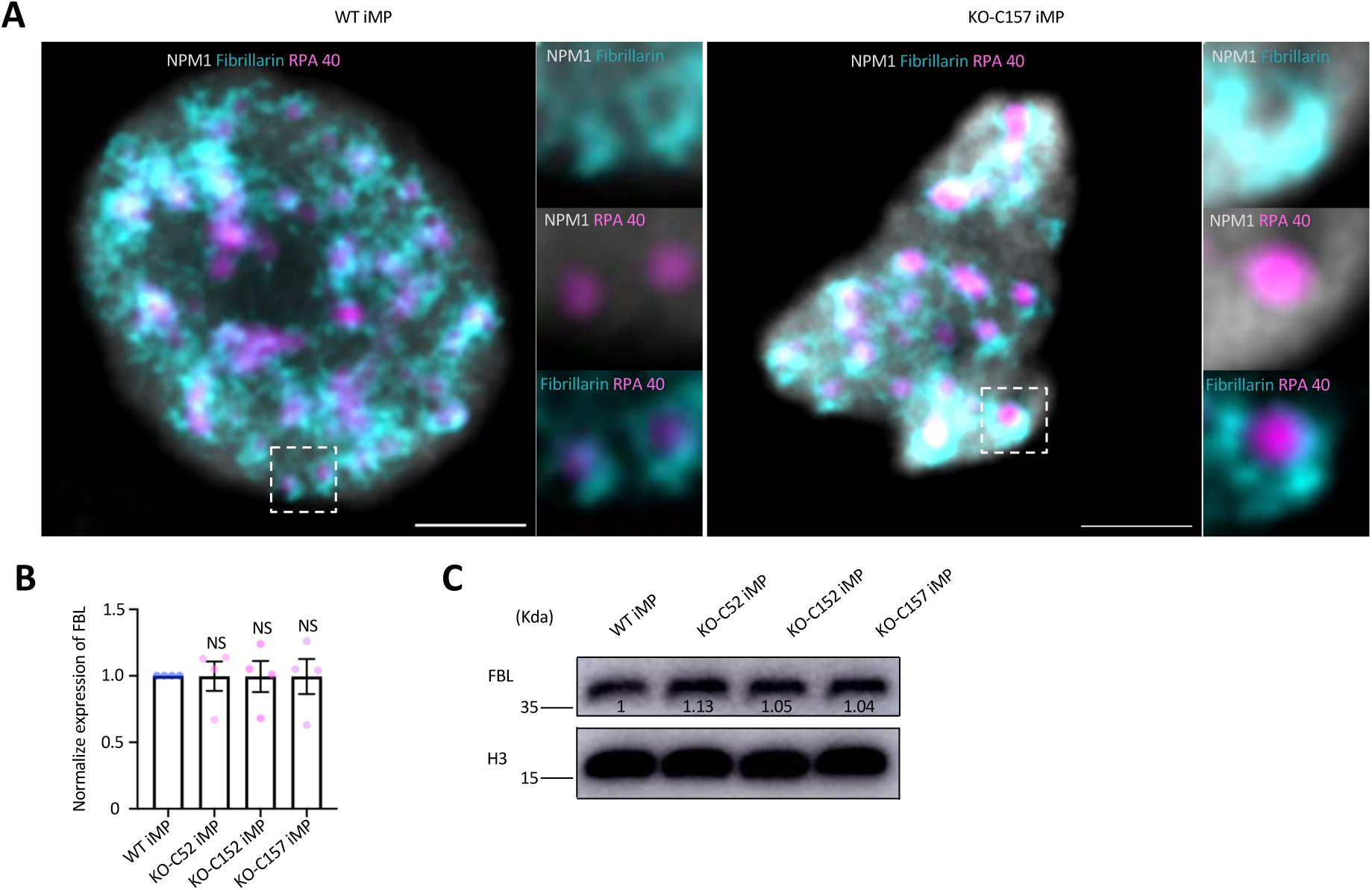
WASP-deficient cells do not alter the sub-localization of the nucleolus and FBL expression, related to Figure 3. A, Representative super-resolution images of the sub-domains of the nucleolus in WT and WASP-KO macrophage cells. NPM1: GC, Fibrillarin: DFC, RPA 40: FC. Bar =1 μm. n = 10. Two biological replicates. B, Quantitative analysis of FBL expression in WT iMP and three WAS KO-iMPs from four biological replicates. C. Representative Western blot analysis of FBL expression in WT iMP and three WAS KO-iMPs. H3 was used as the loading control. The normalized expression value of FBL relative to WT iMPs is indicated by the numbers in the blot. The experiment was independently repeated four times.

**Fig. S3:**
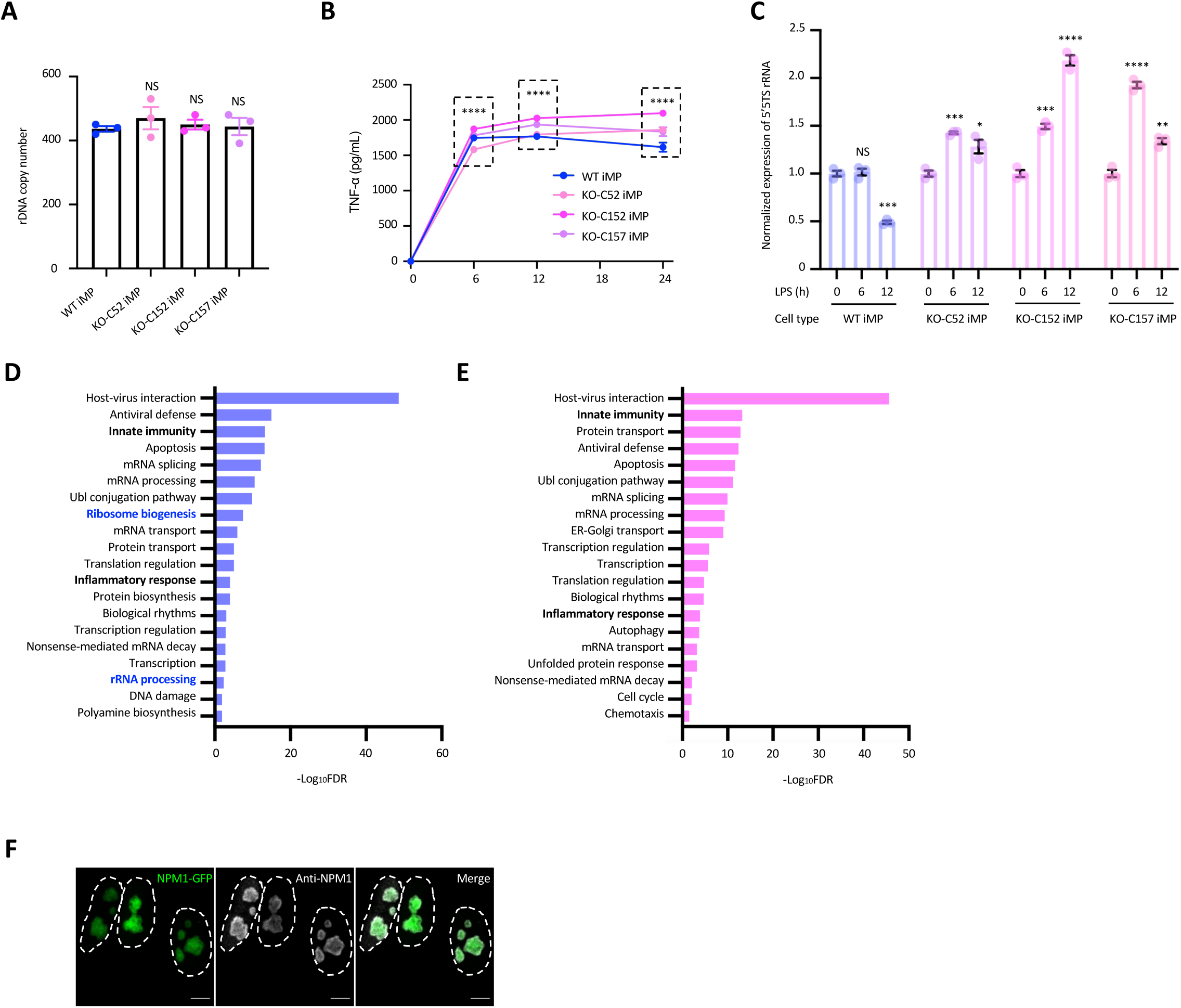
WASP deficiency hampers the ribosomal RNA transcription, related to Figure 4. A, Analysis of 45S rDNA copy number in the indicated iMPs using ddPCR. Data are shown as the mean ± SEM from three biological repeats. Mann-Whitney test. NS, not significant. B, Quantification of TNF-α secretion in WT and WASP-KO macrophages upon stimulation with LPS at different time points. Data are shown as mean ± SEM from two biological repeats. **** p < 0.0001. C, Normalized expression of 5’ETS in WT and WASP-KO macrophages upon stimulation with LPS at the indicated time points analyzed with ddPCR. Data are shown as mean ± SEM from three biological repeats. D, GO analysis of enriched pathways in up-regulated genes in WT iMPs following LPS stimulation. E, GO analysis of enriched pathways in up-regulated genes in WASP-KO iMPs following LPS stimulation. F, Representative images of inducible NPM1-GFP signals and endogenous NPM1 signals. Bar = 5 μm.

---

| ANTIBODIES | SOURCE | IDENTIFIER | DILUTION |
| --- | --- | --- | --- |
| Mouse monoclonal anti-WASP antibody (D1) | Santa Cruz Biotechnology | sc-5300 | WB: 500x |
| Mouse monoclonal anti-WASP antibody (B9) | Santa Cruz Biotechnology | sc-13139 | IF: 100x<br>PLA: 200x<br>WB: 500x |
| RPA194 Antibody (C-1) | Santa Cruz Biotechnology | sc-48385 | IF: 200x<br>WB:1000x |
| RPA40 Antibody (H-6) | Santa Cruz Biotechnology | sc-374443 | WB: 500x |
| RPA40 Antibody (H-6) Alexa Fluor 488 | Santa Cruz Biotechnology | sc-374443 AF488 | IF: 100x |
| Anti-trimethyl-Histone H3 (Lys27) Antibody | Merck | 07-449 | IF: 200x |
| Anti-Mouse-IgG - Atto 647N antibody produced in goat | Merk | 50185-1ML-F | IF: 500x |
| Anti-Rabbit-IgG - Atto 647N antibody produced in goat | Merk | 40839-1ML-F | IF: 500x |
| NPM1 Antibody | Bethyl Laboratories | A302-402A | PLA: 200x |
| Anti-Histone H3 antibody | Abcam | ab10799 | WB: 4000x |
| Anti-Fibrillarin antibody | Abcam | ab5821 | IF/PLA: 1000x<br>WB: 1000x |
| Anti-WASP/Wiskott-Aldrich syndrome protein antibody | Abcam | Ab74904 | IF: 200x |
| NPM1 Monoclonal Antibody (FC-61991) | Thermo Fisher scientific | FC-61991 | IF: 400x<br>WB:1000x |
| IgG (H+L) Cross-Adsorbed Donkey anti-Mouse, HRP | Thermo Fisher scientific | SA1100 | WB: 2000x |
| IgG (H+L) Cross-Adsorbed Donkey anti-Rabbit, HRP | Thermo Fisher scientific | SA1200 | WB: 2000x |
| IgG (H+L) Highly Cross-Adsorbed Donkey anti-Mouse, Alexa Fluor™ 488 | Thermo Fisher scientific | A21202 | IF: 500x |
| IgG (H+L) Highly Cross-Adsorbed Donkey anti-Rabbit, Alexa Fluor™ 488 | Thermo Fisher scientific | A21206 | IF: 500x |
| IgG (H+L) Highly Cross-Adsorbed Donkey anti-Mouse, Alexa Fluor™ 594 | Thermo Fisher scientific | A21203 | IF: 500x |
| IgG (H+L) Highly Cross-Adsorbed Donkey anti-Rabbit, Alexa Fluor™ 594 | Thermo Fisher scientific | A21207 | IF: 500x |
| IgG (H+L) Highly Cross-Adsorbed Donkey anti-Mouse, Alexa Fluor™ 647 | Thermo Fisher scientific | A31571 | IF: 500x |
| IgG (H+L) Highly Cross-Adsorbed Donkey anti-Rabbit, Alexa Fluor™ 647 | Thermo Fisher scientific | A31573 | IF: 500x |

